# Sexually monomorphic wing pigmentation pattern does not contribute to mate choice in *Drosophila guttifera*

**DOI:** 10.1101/2020.05.04.077909

**Authors:** Takuma Niida, Shigeyuki Koshikawa

## Abstract

In many animal groups, sexually dimorphic ornaments are thought to be evolved by intraspecific competition or mate choice. Some researchers pointed out that sexually monomorphic ornaments could also be evolved by mate choice by both sexes or either sex. Many species of fruit fly have sexually monomorphic wing pigmentation. However, involvement of their sexually monomorphic ornaments in mate choice has not been tested. We aimed to examine whether the sexually monomorphic polka-dotted pattern on wings of *Drosophila guttifera* contributes to mate choice. Because *D. guttifera* does not mate in the dark condition at all and courtship sound was not observed, some visual information is likely to be used in mating behaviour. We compared the number of mates between individuals with and without wings, and found that presence of wings influenced mate choice in both sexes. We then compared the number of mates between individuals bearing replaced wings, one group for conspecific *D. guttifera* wings and another group for heterospecific *D. melanogaster* wings with no pigmentation pattern. The effect of conspecific/heterospecific wings was only detected in mate choice by females. By comparison between wild-type and black-painted wings, we found no evidence of contribution of wing pigmentation pattern to mate choice in either sex.

## Introduction

Evolution of sexually dimorphic ornaments has been explained in the theoretical framework of sexual selection, such as Fisher’s runaway process and the handicap principle (Fisher, 1915; Zahavi, 1975). In many animal groups, sexually dimorphic ornaments were suggested to be evolved by intrasexual competition and mate choice (Petrie, Halliday, & Sanders, 1991; Petrie & Halliday, 1994; Andersson, 1982; Ryan, 1985; Theis, Salzburger, & Egger, 2012). Some researchers argued that sexually monomorphic ornaments could evolve before sexually dimorphic ornaments under the evolutionary restriction of sexual dimorphism by genetic correlation or constraint between the two sexes, and that dimorphism evolves by mutations enabling circumvention of the genetic constraint on a long time scale (Lande, 1980). Based on this theory, an organism on the way to acquiring sexual dimorphism can still have a sexually monomorphic ornament. As a counter explanation, the same traits could be involved in mate choices of both sexes, which results in evolution of a sexually monomorphic ornament (Kraaijeveld, Kraaijeveld-Smit, & Komdeur, 2007). King penguins, *Aptenodytes patagonicus*, have sexually monomorphic ornaments, but female mate choice was observed to be stronger than male mate choice (Pincemy, Dobson, & Jouventin, 2009). This suggests that sexually monomorphic ornaments could evolve by a mate choice by either sex.

In *Drosophila* (fruit flies), there are species with various pigmentation patterns on wings (Koshikawa, 2020). Many species of *Drosophila* are known to have male-specific black pigmentation on the anterior-distal part of wings (Kopp & True, 2002). Males of these species appear to display their wings in front of females (Prud’homme et al., 2006). But an effect of wing pigmentation on mate choice was not always observed. Using three species with sexually dimorphic black spots on male wings (*Drosophila suzukii, D. biarmipes* and *D. subpulchrella*), Roy and Gleason (2019) examined whether females prefer males with spots or males without spots, which were made by CO_2_ anesthesia after eclosion. An effect of spots on mating time was not detected in their study. Fuyama (1979) revealed that males without spots as a result of amputation showed lower mating frequency than intact males in *D. suzukii*, when females were kept in constant light to make them less accepting of mating. In *D. biarmipe*s, which has natural polymorphism of wing pigmentation, males with pigmentation on wings showed greater mating success than males without pigmentation (Hegde, Chethan, & Krishna, 2005; Singh & Chatterjee, 1987). The effect of pigmentation was dependent on environmental conditions (Parkash, Lambhod, & Singh, 2013).

Despite having sexually monomorphic pigmentation on their wings, some fruit fly males display their wings in front of females during their courtship. For example, in *Idiomyia grimshawi* (synonym of *Drosophila grimshawi*), adults aggregate in leks and males display with their elaborately pigmented wings (Spieth, 1966; Edwards, Doescher, Kaneshiro, & Yamamoto, 2007). However, the function of this sexually monomorphic pigmentation during courtship has not been studied experimentally. Many species of fruit flies are known to have sexually monomorphic wing pigmentations (Patterson, 1943; Koshikawa, 2020; Werner, Steenwinkel, & Jaenike, 2018), but no functional testing of these monomorphic traits has been reported.

*Drosophila guttifera* has been used as a research model for elucidating pigmentation pattern formation (Werner, Koshikawa, Williams, & Carroll, 2010; Koshikawa et al., 2015; Koshikawa, Fukutomi, & Matsumoto, 2017; Fukutomi, Matsumoto, Agata, Funayama, & Koshikawa, 2017; Fukutomi, Kondo, Toyoda, Shigenobu, & Koshikawa, 2020). This species has a sexually monomorphic polka-dotted pattern on its wings, but the function of the pattern has not been explored. Because *D. guttifera* does not mate in a dark condition at all, visual information is likely to be used in mating (Grossfield, 1966). In addition, courtship sound, such as wing vibration, was not observed in this species (Spieth, 1952; Grossfield, 1977; Wen & Li, 2011).

The purpose of this study was to clarify whether the sexually monomorphic pigmentation pattern in *D. guttifera* contributes to mate choice. We examined the effect of the presence of wings and the polka-dotted pattern on wings on mate choice of both sexes, by conducting mating experiments with cutting and replacing of wings.

## Materials and methods

### Fly

*Drosophila guttifera* inhabits North America and is related to the *quinaria* group (Chialvo, White, Reed, & Dyer, 2019; Izumitani, Kusaka, Koshikawa, Toda, & Katoh, 2016). In this study, adults of the wild type *D. guttifera* (stock number 15130-1971.10, provided by the Drosophila Species Stock Center at the University of California, San Diego) kept in our laboratory were used. In addition to intact adults (“wild type”), adults whose wings were cut within 24 hours after eclosion (“no wing”) and adults whose wings were painted black (“black wing”) were prepared. Also, adults whose wings were replaced by *D. guttifera* wings (“*guttifera* wing”), wings with an incomplete pigmentation pattern (“incomplete pattern”), wings of *Drosophila melanogaster* Oregon-R (“*melanogaster* wing”), and black-painted wings of *D. melanogaster* (“*melanogaster* black wing”) were prepared. These flies were used in mate choice experiments (Figure 1). The strain with an incomplete pigmentation pattern has a recessive mutation, but the genetic lesion of this mutation is unknown (Wataru Yamamoto, personal communication). Wings were replaced using a cyanoacrylate adhesive (Aron Alpha, Konishi Co., Ltd., Japan). To produce black wing flies, their entire wings were painted with a black paint marker (Mckie, Zebra Co., Ltd., Japan). All individuals were reared with standard cornmeal/sugar/yeast/agar food under a photoperiod of 12:12 (light:dark) at 25°C (Fukutomi, Matsumoto, Funayama, & Koshikawa, 2018).

**Figure 1.**
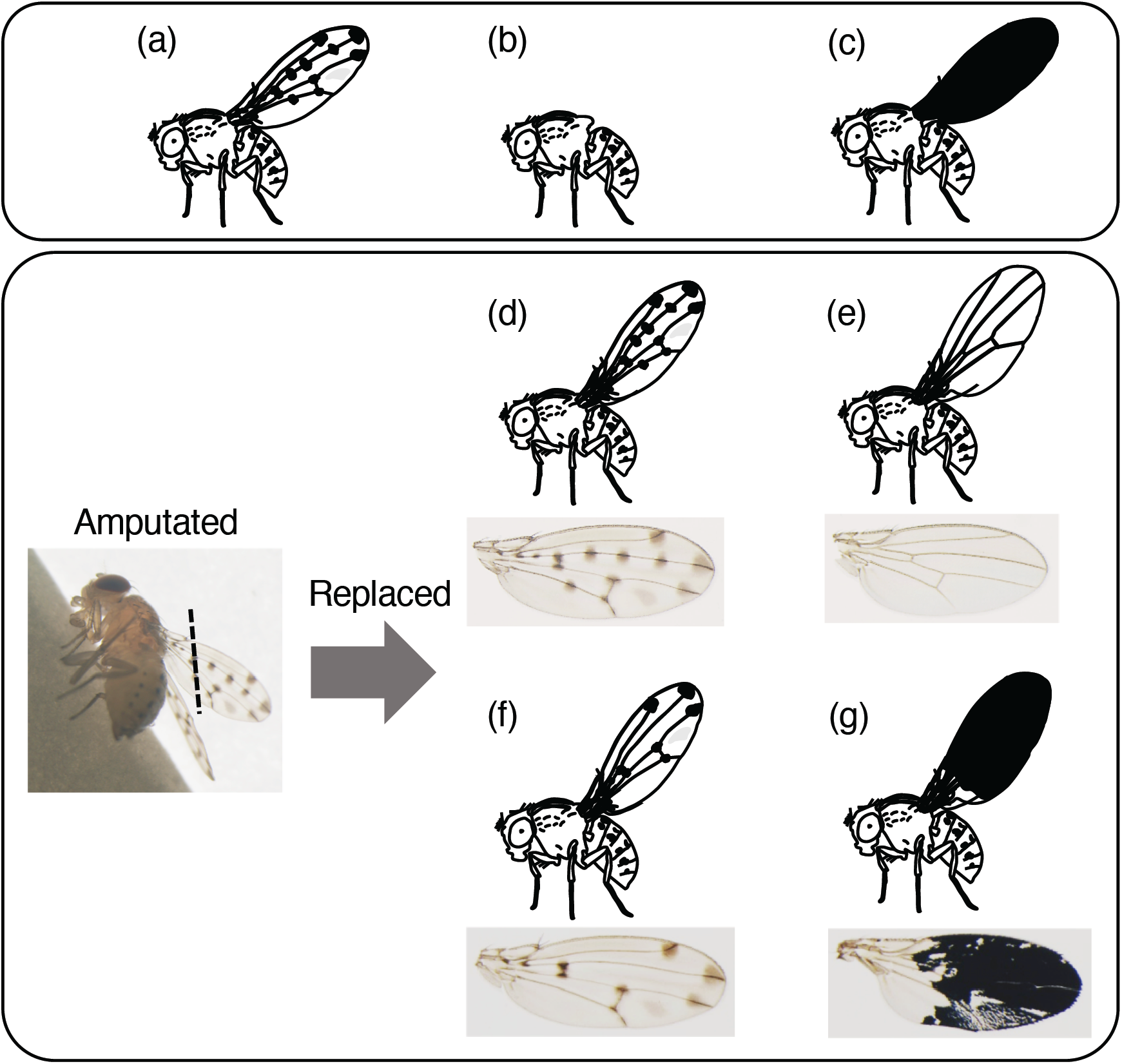
Flies used in mating experiments. (a) A “wild type” adult. (b) A “no wing” adult, whose wings were removed. (c) A “black wing” adult, whose wings were painted black. (d) A “*guttifera* wing” adult, whose wings were replaced with *D. guttifera* wings. (e) A “*melanogaster* wing” adult, whose wings were replaced with *D. melanogaster* wings. (f) An “incomplete pattern” adult, whose wings were replaced with wings without pigmentation spots around the campaniform sensilla. (g) A “*melanogaster* black wing” adult, whose wings were replaced with *D. melanogaster* wings painted black

### Sexual maturity after eclosion

To prepare virgin individuals for mating experiments, we had to confirm that adults were not sexually mature within 24 hours after eclosion. Firstly, males and females were separated into different vials within 4 hours after eclosion. Some vials were kept 7– 10 days after eclosion. Four females 7–10 days after eclosion and four males 4 hours after eclosion were introduced into one vial. Likewise, four males 7–10 days after eclosion and four females 4 hours after eclosion were introduced into one vial. In a control group, four females and four males 7–10 days after eclosion were introduced into one vial. These three groups of vials were kept for 24 hours, and then adults in the vials were removed. The number of larvae in the vials was counted 5–7 days after adults had been removed.

### Recording of mating behaviour

All mate choice experiments were conducted in chambers made of acrylic resin (10 mm in diameter and 5 mm in height). A small piece of fly food was placed in each chamber to prevent drying of flies during experiments. The chambers were put on a piece of white paper, and then adults used in experiments were introduced into the chambers. The chambers were then illuminated with a halogen lamp (PICL-NEX, Nippon P·I, Co., Ltd.) and recorded with a web camera (C615n, Logicool, Co., Ltd., Japan) or a digital camera (HDR-PJ590V, Sony, Japan) continuously for 3 hours. The start time of the recording was between 90 and 150 minutes after light phase of the original rearing condition.

### Male mate choice experiments

Males and females were removed from the vials in which they had been reared, and were separated, within 24 hours after eclosion, and then male mate choice experiments were conducted 5–13 days after eclosion. To examine the preference toward wings, we observed whether males mated with “wild type” females or “no wing” females. Next, to examine the preference toward the polka-dotted pattern, we observed whether males mated with “*guttifera* wing” females or “*melanogaster* wing” females. If mating was observed, we recorded the copulation duration.

### Female mate choice experiments

Males and females were removed from the vials in which they had been reared within 24 hours after eclosion, and were separated, and then female mate choice experiments were conducted 5–13 days after eclosion. To examine the preference toward wings, we observed whether females mated with “wild type” males or “no wing” males. Next, to examine the preference toward the polka-dotted pattern, we observed whether females mated with “*guttifera* wing” males or “*melanogaster* wing” males. To examine the preference toward the precise number of polka dots, we observed whether females mated with “*guttifera* wing” males or “incomplete pattern” males. Also, to examine the preference toward dark color of the entire wings, we observed whether females mated with “wild type” males or “black wing” males. Finally, we observed whether females mated with “*melanogaster* wing” males or “*melanogaster* black wing” males. If mating was observed, we recorded the copulation duration.

### Statistical analysis

All statistical analyses were performed using R ver. 3.5.2 (R Core Team, 2018). When we constructed multiple models, the model with the lowest AIC (Akaike’s information criterion) was chosen. In male mate choice experiments, generalized linear models (GLMs) were used to investigate the influence of male and female age, the influence of the mating combination between the sexes, and hours after females had received replaced wings, on the occurrence of mating. The occurrence of mating was treated as a response variable assuming a binominal distribution. Male and female age, mating combination between the sexes, and hours after they had received replaced wings were treated as explanatory variables.

The kinds of mating were classified into four types—the first mating between one male and one female, the second mating between them, the first mating between one male and the other female, and the second mating between them. GLMs were used to examine the influence of male age, female age, and mating order on which females mated with one male. Which females mated with one male was treated as a response variable assuming a binominal distribution. Male age, female age and mating order were treated as explanatory variables. Multiple comparisons of copulation durations were performed using the Tukey-Kramer test.

In female mate choice experiments, GLMs were used to examine the influence of male age, female age, mating combination between the sexes, and hours after males had received replaced wings on the occurrence of mating. The occurrence of mating was treated as a response variable assuming a binominal distribution. Male age, female age, mating combination between the sexes, and hours after the males had received replaced wings were treated as explanatory variables. The kinds of mating were classified into two types—mating between one female and one male, and mating between one female and the other type of male. Copulation durations were compared using Student’s *t*-test.

## Results

### Sexual immaturity at 24 hours after eclosion

Four males 7–10 days after eclosion and four females 4 hours after eclosion were introduced into one vial and were removed from the vial after 24 hours. After 5–7 days, no larvae were observed in the vial (Table 1). Also, no larvae were observed in vials into which four females 7–10 days after eclosion and four males 4 hours after eclosion were introduced and then removed. In contrast, the number of larvae was 15.8 ± 11.0 (mean ± SD [standard deviation]) in vials into which four males and four females 7–10 days after eclosion were introduced and then removed.

**Table 1.** Sexual maturity test. The number of larvae in three groups (mean ± SD [standard deviation])

|  | 4 females 7–10 days<br>after eclosion<br>+<br>4 males 4 hours<br>after eclosion (n=5) | 4 males 7–10 days<br>after eclosion<br>+<br>4 females 4 hours<br>after eclosion (n=6) | 4 females and 4<br>males 7–10 days<br>after eclosion (n=4) |
| --- | --- | --- | --- |
| Number of larvae | 0 | 0 | 15.8±11.0 |

We concluded that neither males nor females become sexually mature within 24 hours after eclosion. Therefore, in the following experiments, the adults within 24 hours after eclosion were collected as virgins. We kept virgins for more than 5 days and used them in mate choice experiments.

### Male preference

In experiments to examine male preference, some males mated with both of the two females. Such matings were recorded separately as the first mating and the second mating. Males never mated twice with the same female in our observations.

We conducted 51 experiments to examine preference for the presence of wings (Figure 2a). In the first matings, males mated with “wild type” females 25 times and with “no wing” females 10 times. In the second matings, males mated with “wild type” females four times and with “no wing” females 11 times. Thus, males mated with “wild type” females at a higher rate than with “no wing” females (p=0.0244, GLM). Moreover, males mated at a higher rate with “wild type” females at the first mating, and at a higher rate with “no wing” females at the second mating (p=0.000682, GLM). The copulation duration of the first mating between males and “wild type” females was 475.6 ± 42.3 seconds (mean ± SD) and that of the second mating between them was 438.8 ± 76.4 seconds (Figure 2b). The copulation duration of the first mating between males and “no wing” females was 449.4 ± 63.4 seconds and that of the second mating between them was 424.5 ± 44.9 seconds. The copulation duration of the first mating between males and “wild type” females was significantly longer than the copulation duration of the second mating between males and “no wing” females (p=0.0323, Tukey-Kramer test). In summary, males were found to prefer females with wings to females without wings.

**Figure 2.**
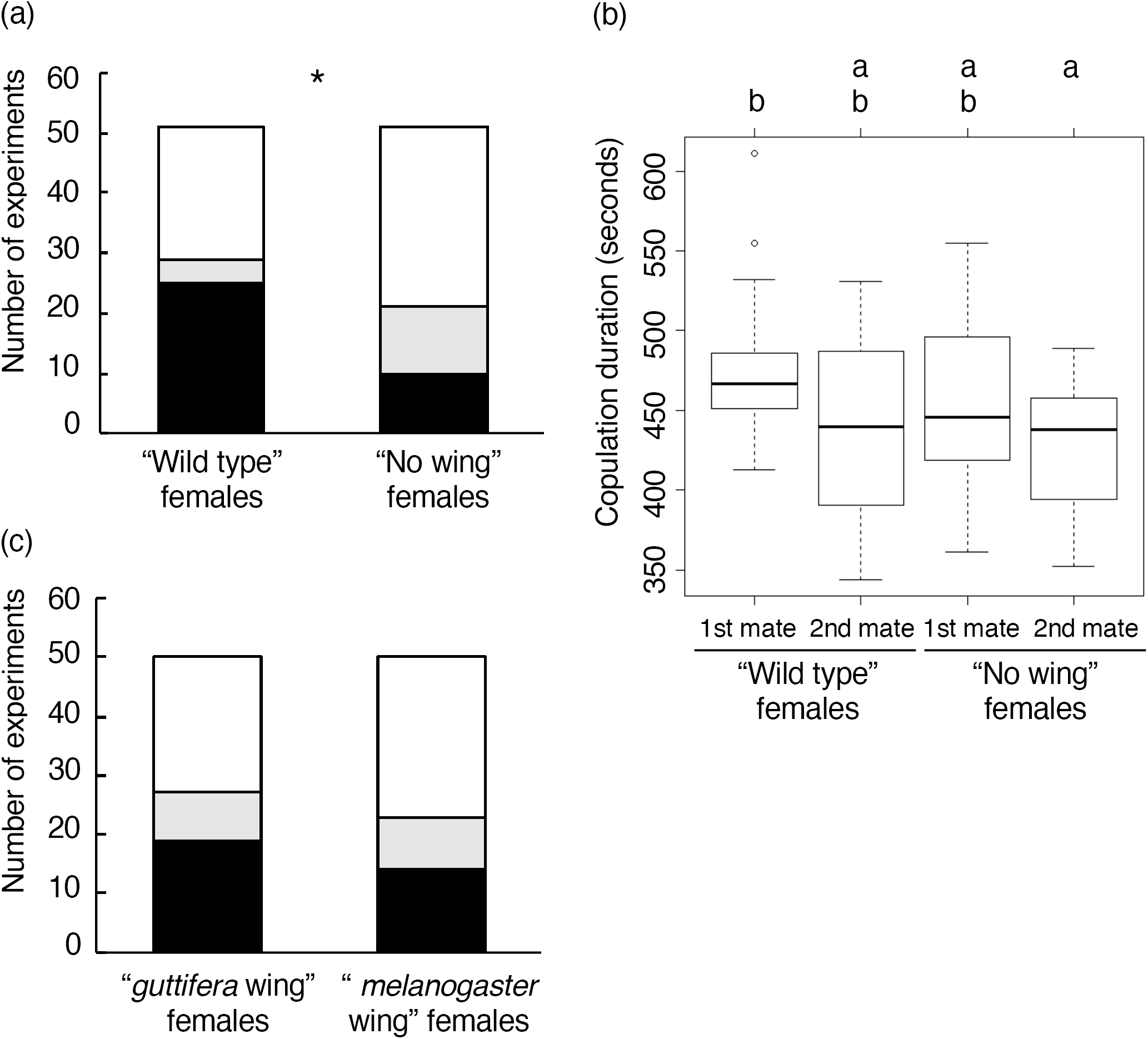
Male mate choice experiments. (a) Preference for females with/without wings. Black areas of the graph represent the first mates. Grey areas represent the second mates. White areas represent no mates. (b) Copulation durations between males and “wild type” females, males and “no wing” females in the first and second matings, respectively. (c) Preference for females with “*guttifera wing*” or “*melanogaster wing*”. Asterisk (*) indicates significance at p <0.05

We conducted 50 experiments to examine the preference for the polka-dotted pattern (Figure 2c). In the first matings, males mated with “*guttifera* wing” females 19 times and “*melanogaster* wing” females 14 times. In the second matings, males mated with “*guttifera* wing” females 8 times and *D. melanogaster* wing females 9 times. No significant difference was seen between the rate of mating of males with “*guttifera* wing” wing females, and that of males with “*melanogaster* wing” females (p=0.544, GLM). The copulation duration of the first mating between males and “*guttifera* wing” females was 536.4 ± 70.5 seconds, and that of the second mating between them was 480.5 ± 182.8 seconds. The copulation duration of the first mating between males and “*melanogaster* wing” females was 517.2 ± 48.2 seconds and that of the second mating between them was 474.5 ± 35.3 seconds. No significant difference was observed in these copulation durations (p=0.32, One-way ANOVA). In summary, there was no tendency for males to prefer “*guttifera* wing” females to “*melanogaster* wing” females.

### Female preference

In experiments to examine female preference, all females mated with only one of the males. Females never mated twice with the same male in our observations.

We conducted 51 experiments to examine preference for wings (Figure 3a). Females mated with “wild type” males 27 times and with “no wing” males 15 times. Females mated with “wild type” males more frequently than with “no wing” males (p=0.0158, GLM). Copulation duration between females and “wild type” males was 464.4 ± 58.7 seconds. Copulation duration between females and “no wing” males was 494.8 ± 49.1 seconds. No significant difference was observed between these copulation durations (p=0.1287, t-test). In summary, females were found to prefer males with wings to males without wings.

**Figure 3.**
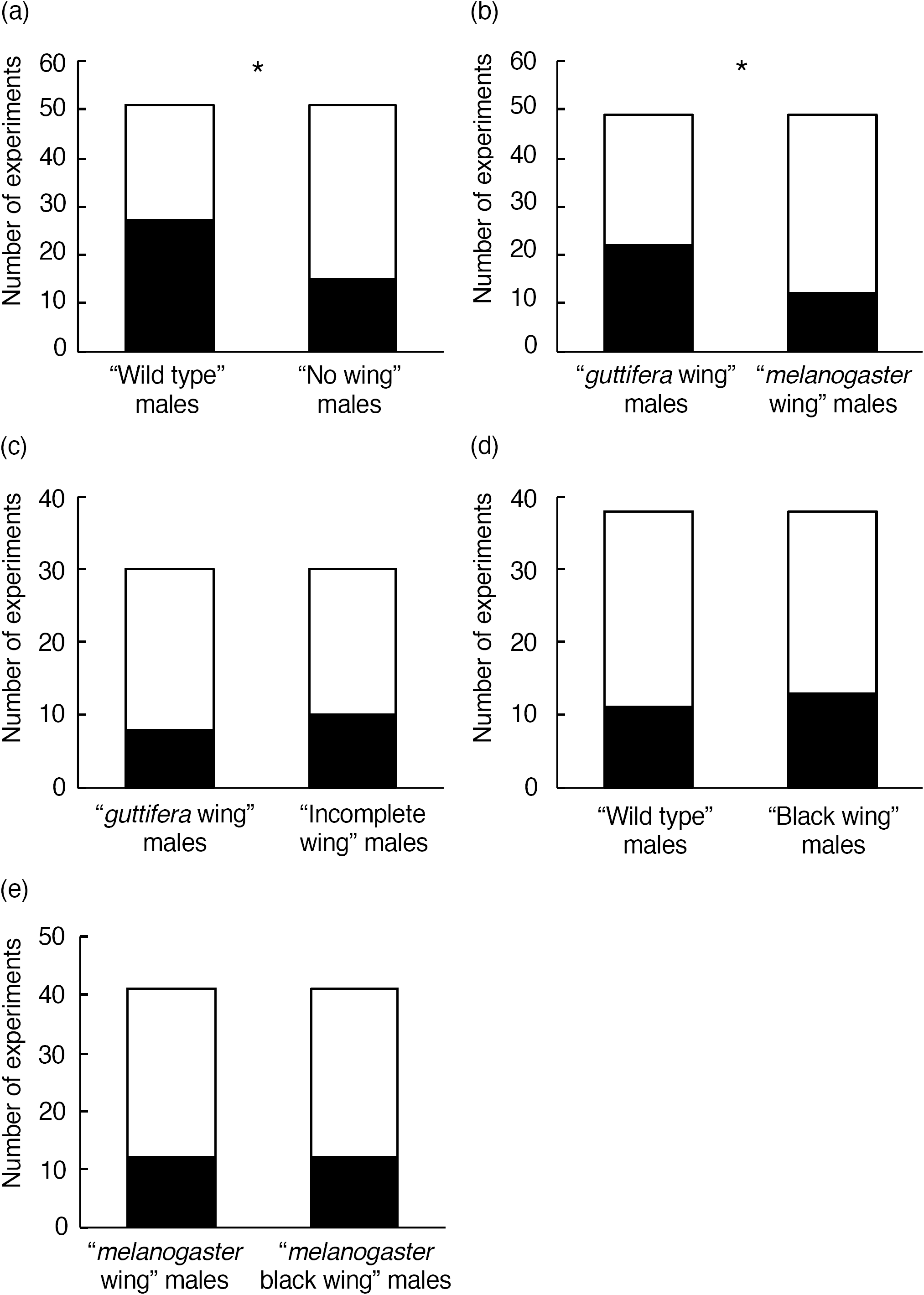
Female mate choice experiments. (a) Preference for males with/without wings. Black areas of the graph represent the first mates. White areas represent no mates. (b) Preference for males with “*guttifera wing*” or “*melanogaster wing*”. (c) Preference for males with “*guttifera wing*” or “incomplete pattern”. (d) Preference for males of “wild type” or with “black wing”. (e) Preference for males with “*melanogaster* wing” or “*melanogaster* black wing”. Asterisks (*) indicate significance at p <0.05

We conducted 49 experiments to examine preference for polka-dotted *D. guttifera* wings (Figure 3b). Females mated with “*guttifera* wing” males 22 times and “*melanogaster* wing” males 12 times. Females mated with “*guttifera* wing” males significantly more frequently than with “*melanogaster* wing” males (p=0.0361, GLM). Copulation duration between females and “*guttifera* wing” wing males was 530.9 ± 74.5 seconds. Copulation duration between females and “*melanogaster* wing” males was 542.5 ± 50.7 seconds. No significant difference was observed between these copulation durations (p=0.6429, *t*-test). In summary, females were found to prefer “*guttifera* wing” males to “*melanogaster* wing” males. In these experiments, however, we could not distinguish whether females preferred conspecific wings or the polka-dotted pattern. For this reason, we performed additional experiments using conspecific but differently patterned wings.

In these additional experiments, we conducted 30 experiments to examine preference for different pigmentation patterns (Figure 3c). Females mated with “*guttifera* wing” males eight times and “incomplete pattern” males 10 times. No significant difference was observed between mating times in females and “*guttifera* wing” males, and those in females and “incomplete pattern” males (p=0.572, GLM). Copulation duration between females and “*guttifera* wing” males was 692.8 ± 71.4 seconds. Copulation duration between females and “incomplete pattern” males was 660.6 ± 155.6 seconds. No significant difference was observed between these copulation durations (p=0.5977, *t*-test). In summary, no tendency was found for females to prefer different pigmentation patterns.

We then considered the possibility that females might prefer darkness of male wings, not a particular pattern. We conducted 38 experiments to examine preference for darkness of entire wings (Figure 3d). Females mated with “wild type” males 11 times and “black wing” males 13 times. No significant difference was observed between mating times in females and “wild type” males, and those in females and “black wing” males (p=0.6219, GLM). Copulation duration between females and “wild type” males was 746.7 ± 136.1 seconds. Copulation duration between females and “black wing” males was 736.5 ± 80.4 seconds. No significant difference was observed between these copulation durations (p=0.8207, *t*-test). In summary, no tendency was found for females to prefer darkness of entire wings.

Females thus did not show a preference regarding different pigmentation patterns or regarding darkness of the entire wings. Females thus did not seem to care about the details of wing pigmentation. Finally, we examined whether females preferentially mated with “*melanogaster* wing” males or “*melanogaster* black wing” males (Figure 3e). Females mated with “*melanogaster* wing” males 12 times and “*melanogaster* black wing” males 12 times. In addition, no significant difference was observed between mating time in females and “*melanogaster* wing” males, and that in females and “*melanogaster* black wing” males (p=0.9861, GLM). Copulation duration between females and “*melanogaster* wing” males was 680.8 ± 216.6 seconds. Copulation duration between females and “*melanogaster* black wing” males was 811.2 ± 68.0 seconds. No significant difference was observed between these copulation durations (p=0.0592, t-test). Taking these results altogether, we concluded that females prefer *D. guttifera* wings to *D. melanogaster* wings, but we found no preference for black-painted compared to unpainted *D. melanogaster* wings. Therefore, the difference between the preference for *D. guttifera* wings and *D. melanogaster* wings does not seem to be caused by wing color or pattern, but by some other wing characteristic(s), such as wing shape or smell.

## Discussion

### Preference for wings

Previous studies showed that wing damage can influence reproductive success, survival, mortality and flight performance in various insects, such as Odonata (Combes, Crall, & Mukherjee, 2010), Hymenoptera (Cartar, 1992) and Lepidoptera (Jantzen & Eisner, 2008). The present study found that both males and females mate more with intact wild-type adults than adults without wings. This suggests that wing damage could influence the preference for mates in *D. guttifera*. Because previous studies showed that *D. guttifera* males do not perform courtship with wing vibration (Spieth, 1952; Grossfield, 1977; Wen & Li, 2011), mate choice by females does not seem to require male wing motion. Instead, females may recognize male wing smell or shape as a survival value and use these features to choose mates.

Similarly, males might use the presence of wings to assess the survival value of females. Male mate choice has been reported in multiple *Drosophila* species (Bonduriansky, 2001; Byrne & Rice, 2006). Another potential explanation for female wing function is related to females’ behaviour during a courtship sequence. At the moment of mate acceptance, females spread their wings widely, and this motion is likely to work as a visual signal of acceptance (Spieth, 1952; Grossfield, 1966). This could be a reason why females without wings had a lower rate of mating than intact females.

### Preference for conspecific wings

In male mate choice experiments, no significant difference was seen between mating times in males and “*guttifera* wing” females, and those in males and “*melanogaster* wing” females. But in female mate choice experiments, females were found to mate more with “*guttifera* wing” males than “*melanogaster* wing” males. Because female mate choice generally is stronger and more common than male mate choice (Bonduriansky, 2001), these results would not be surprising.

One of reasons for the sexual difference in the preference for conspecific wings could be the biased operational sex ratio. During 3 hours of recording, males mated with multiple females, but females mated with only one male. A difference in mating receptivity between the sexes may influence operational sex ratio, which is the number of sexually active males and females (Markow, 1988). In *D. guttifera*, female mating receptivity is likely to be lower than male mating receptivity. Consequently, females could develop mate choice toward males (Clutton-Brock & Vincent, 1991), and conspecific wings, of course, could be one of the criteria to prefer.

### Preference for the polka-dotted pattern

In male mate choice experiments, no significant difference was observed between mating times in males and “*guttifera* wing” females, and those in males and “*melanogaster* wing” females. *D. melanogaster* does not have a pigmentation pattern on its wings. Therefore, this result suggested that males do not have a preference for the polka-dotted pattern or any other trait of conspecific wings.

In female mate choice experiments, females mated more with “*guttifera* wing” males than “*melanogaster* wing” males. This result suggested two possibilities—females have a preference for the polka dotted-pattern or a preference for other trait(s) of conspecific wings. Further experiments showed that females did not have a preference for the details of the polka-dotted pattern or for darkness of the entire wings.

These experiments suggest that the sexually monomorphic polka-dotted pattern on wings does not contribute to mate choice of either of the sexes in *D. guttifera*. To our knowledge, this is the first study to examine the contribution of sexually monomorphic ornaments in *Drosophila* to mate choice. We cannot rule out, however, the possibility of a weak preference for the polka-dotted pattern, which was not statistically significant in this study, but is strong enough to be a selective pressure to fix the pattern in the population. So far, the contribution of sexually monomorphic ornaments to mate choice is known in king penguins (Pincemy et al., 2009; Nolan et al., 2010). In the future, it should be verified what condition(s) enables sexually monomorphic ornaments to contribute to mate choice, using species of various groups.

A contribution to mate choice was not detected in this study, and the function of the polka-dotted pattern on wings of *D. guttifera* is still unclear. Considering the extremely sophisticated gene regulation that acts to form the polka-dotted pattern (Koshikawa, 2020; Fukutomi et al., 2020), it is difficult to think the pattern randomly evolved as neutral trait without adaptive significance. Future research is needed to reveal what adaptive significance(s) the polka-dotted pattern has in *D. guttifera*.

## Acknowledgements

We thank Kiyohito Yoshida and Wataru Yamamoto for technical help; Yuichi Fukutomi, Tomohiro Yanone, Wataru Yamamoto, Masato Koseki and Namiho Saito for discussions; and Elizabeth Nakajima for English editing. We also thank Kyosuke Okawara and Manabu Kamimura for their advice in the early stage of this study. This study was supported by KAKENHI (17K19427 and 18H02486).

## Conflict of interest

The authors declare that they have no conflict of interest.

## Notes

### Competing Interest Statement

The authors have declared no competing interest.

